# Negative Autoregulation Promotes the Evolution of Strong Environmental Switching

**DOI:** 10.64898/2026.06.30.735141

**Authors:** Pouria Dasmeh, Gopinath Chattopadhyay, Diego Pesce, Cauã Westmann, Andreas Wagner

## Abstract

Living systems rely on gene regulatory circuits to respond to environmental change. Such circuits often act as molecular switches: OFF in the absence of a cue, and ON in its presence. We do not know how a circuit’s regulatory architecture affect its ability to evolve such responsiveness. In nature, a very frequent and simple regulatory architecture involves a transcriptional regulator that negatively regulates its own expression. To study how such autoregulation affects adaptive evolution, we engineered *E. coli* circuits in which a target gene is regulated by the repressor TetR with (A−) or without (A0) negative autoregulation of TetR. We evolved TetR in both architectures toward responsiveness to a novel environment, embodied by a novel inducer of TetR. Early during their evolution, TetR circuits with negative autoregulation evolved stronger environmental switching. A combination of high throughput DNA sequencing, protein engineering, and biophysical modeling showed why. Only A-circuits favored TetR alleles that interact strongly with both the inducer and DNA. Such alleles combine strong repression caused by strong DNA binding with strong derepression caused by strong inducer binding, the defining property of a strong environmental response. Our biophysical model shows that negative autoregulation helps to create this regulatory regime. As a result, only A^−^ circuits favor alleles that create strong molecular switches. Altogether, our work shows that even the simplest form of gene regulation can change the topography of a fitness landscape, and enable new modes of evolutionary change.

## Introduction

The processes of life are regulated at various scales within an organism, from that of individual molecules to that of cells, tissues, and organs. Much of this regulation involves circuits of regulatory molecules. An especially important class of such circuits are gene regulatory circuits, in which proteins called transcription factors bind specific regulatory DNA sequences near genes and activate or repress their expression^1,2^. Some of the regulated genes may themselves encode transcription factors^3^. Regulatory circuits are crucial to help organisms respond to changing environments. They tune levels of key proteins so that cells can cope with incessantly changing amounts of nutrients and stressors.

The response of a gene regulatory network to the environment often resembles the behavior of a molecular switch. In the absence of an environmental cue, an efficient circuit remains in the OFF state, minimizing unnecessary gene expression. In the presence of the cue, the same circuit flips to the ON state, driving strong expression of the relevant genes. Such switch-like responsiveness can play a crucial role in the evolutionary adaptation of organisms to new environments^4–8^. A classic example is the *E. coli* lac operon, which switches between OFF and ON states depending on the availability of lactose. Evolution experiments show that mutations in this circuit can tune the activation threshold of gene expression, thereby optimizing fitness in changing sugar environments^9,10^. Another example is the two-component regulatory system PhoP/Q, which enables bacteria to survive antimicrobial stress. The sensor kinase PhoQ detects environmental stressors such as antimicrobial peptides, and phosphorylates the transcription factor PhoP, which then activates genes involved in resistance, including efflux pumps^11,12^. In pathogenic bacteria, the two-component system PhoP/Q has undergone adaptive changes that alter the sensitivity of its sensor kinase PhoQ to host-derived antimicrobial peptides, enabling *Salmonella* to fine-tune the induction of resistance genes during infection^13^. These examples underscore a general principle: Natural selection acts not only on protein function but also on regulatory architectures that shape expression dynamics across environments^14^.

To ask whether some forms of gene regulation can convey better environmental switching than others is important for two reason. First, it can help to understand the adaptive evolution of organisms in fluctuating environments and its limits. Second, it can help to design synthetic circuits with robust ON/OFF control. We addressed this question by evolving two distinct regulatory architectures of a regulatory switch, one with and one without negative autoregulation. Our switch is TetR (tetracycline resistance transcriptional repressor), a well-characterized bacterial transcription factor central to the regulation of tetracycline resistance^15^. In the absence of its inducer tetracycline, TetR binds to operator DNA sequences and represses transcription of downstream genes, most notably those encoding efflux pumps^16^. When tetracycline is present, TetR binds the inducer with high affinity, undergoes a conformational change, and detaches from DNA, thereby permitting gene expression^15,17^.

We focus on negative autoregulation, where a transcription factor represses its own expression, because it is one of the most prevalent modes of gene regulation. Negative autoregulation is involved in regulating ∼40% of transcription factors in *E. coli*^18^, and it is widespread in other bacteria, as well as in eukaryotes^19,20^. Circuits with negative autoregulation can provide multiple physiological benefits: they reduce stochastic noise in gene expression^21^, enable more precise control of protein levels^22^, and accelerate response times to environmental changes^23^. By evolving TetR circuits with and without negative autoregulation, we asked whether autoregulation also provides evolutionay benefits. That is, does it facilitate the evolutionary adaptation of a molecular switch to novel environmental cues?

To study the evolution of simple TetR circuits with and without negative autoregulation, we engineered two regulatory circuits. In the first, TetR represses both its own expression and that of a target gene encoding green fluorescent protein (GFP). In the second, TetR represses only the target gene but not itself. We refer to these circuits as the A⁻ and the A□ circuit. We evolved both to regulate GFP expression in response to the novel inducer incyclinide (I_new_). We then asked in which of these circuits TetR evolves to confer greater responsiveness to the environment. We define this responsiveness as the fold-change in the expression of TetR upon addition of the new inducer I_new_. Strong responsiveness has to meet two requirements. First, TetR must interact strongly with DNA to ensure tight repression in the absence of inducer. Second, it must also strongly interact with the inducer to enable efficient derepression when the inducer is present.

We observed that early during directed evolution, negative autoregulation indeed allows A^−^ circuits to evolve greater responsiveness to the environment (the inducer I_new_) than AO circuits. In addition, TetR variants rise to high frequency that increase the interaction strength of TetR with DNA only in A^−^ circuits. A simple biophysical evolutionary model of TetR repression helps to explain why.

## Results

### Engineering the genetic circuits with negative autoregulation

We first engineered the autoregulatory circuit A^−^, in which the TetR dimer binds to the operator tetO and represses its own transcription (Figure 1A). We accomplished this by placing a single TetR operator binding site after the −35 and −10 elements of the TetR promoter, and before the start codon of the TetR protein coding sequence (Figure 1A). In addition, we also engineered the same system but without negative autoregulation (A^0^) by replacing the tetO sequence with a scrambled DNA sequence that can no longer convey negative autoregulation (Materials and methods, Supplementary methods 4). This procedure ensures that the DNA molecules encoding the two circuits have the same length and GC content. To monitor the activity of the TetR repressor, we used a green fluorescent protein (GFP) under the control of the TetR repressor. In the absence of inducer, TetR represses the expression of GFP, causing reduced fluorescence, which we detected by flow cytometry (Methods). The presence of inducer inactivates TetR, which leads to increased GFP expression.

**Figure 1.**
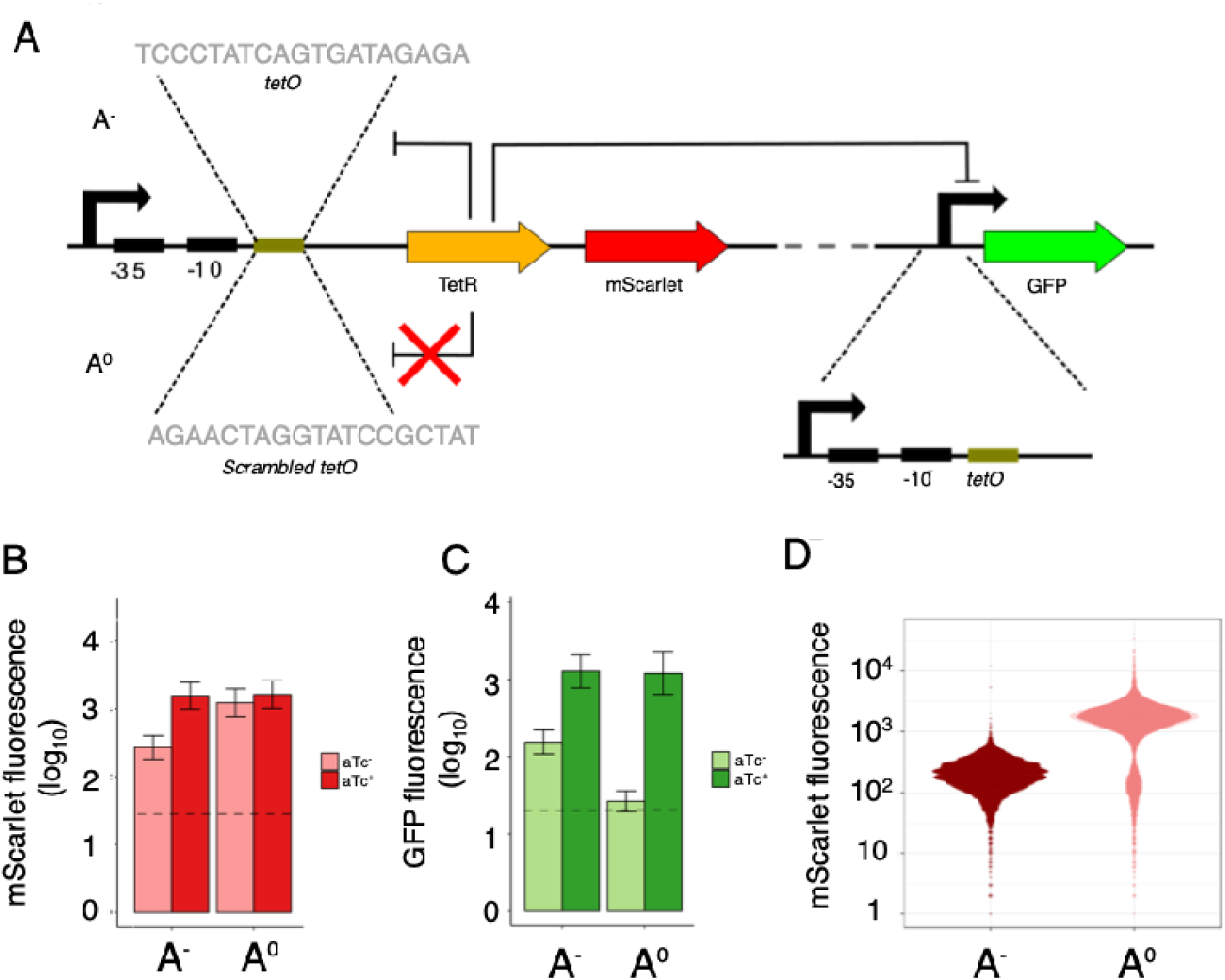
Experimental design of the adaptive evolution experiment. A) Schematic representation of the two circuits constructed for this study: TetR with negative autoregulation (A^−^ circuit) and TetR without negative autoregulation (A^0^ circuit), where the tetO binding site was replaced with a scrambled sequence. B, C) Expression levels of mScarlet (panel B) and GFP (panel C) in the A^−^ and A^0^ circuits, shown as mean fluorescence intensity on a logarithmic scale (base 10). Light bars represent the repressed state (aTc−), while dark bars represent the induced state (aTc+). Dashed lines indicate the background level of cellular autofluorescence. Error bars correspond to one standard error of the mean (SEM). D) Density distribution of mScarlet fluorescence in the A^−^ (dark red) and A^0^ (light red) circuits based on measurements from 10,169 and 11,367 cells, respectively. mScarlet fluorescence serves as a proxy for TetR expression, demonstrating that negative autoregulation reduces both the mean and variance of TetR expression.

In our system, GFP fluorescence serves as a direct readout of the circuit’s response to inducer binding. This response integrates the total derepression activity of the circuit, which includes both changes in TetR binding to the operator and TetR expression level. Both factors are physiologically relevant for regulation of target genes by the circuit^15,24,25^. To monitor TetR expression levels directly, we coupled the TetR gene to a gene encoding the red fluorescent protein mScarlet. This coupling allowed us to measure mScarlet fluorescence as a proxy for TetR expression, since the two genes are expressed from the same transcript. Thus, mScarlet fluorescence provides a quantitative readout of how TetR expression changes under changing conditions.

After constructing both circuits, we first verified their functionality experimentally. In the absence of an inducer, the A^−^ circuit should express less TetR than the A^0^ circuit, because negative autoregulation inherently reduces TetR expression. Our experiments confirm this prediction (Figure 1B). Conversely, when the inducer is present, this difference is expected to vanish because the inducer inactivates TetR. Consistent with this expectation, our experiments showed that the A^−^ and A^0^ circuits express similar levels of TetR in the presence of the inducer (Figure 1B). In addition, because of the lower expression of TetR in the A^−^ circuit when inducer is absent, one would expect that GFP is less strongly repressed in this circuit. Our experiments confirm this prediction as well (Figure 1C).

We also analyzed fluorescence variation within isogenic *E. coli* populations carrying either circuit, because negative autoregulation is known to reduce expression variation caused by gene expression noise^21^. This reduced variation should manifest in less variation in TetR expression and its proxy mScarlet fluorescence. Indeed, the coefficient of variation (CV) of mScarlet fluorescence was significantly lower in A^−^ populations than in A^0^ population (*p* = 0.008, Feltz and Miller asymptotic test for equality of CVs, *p* = 0.0075, modified signed-likelihood ratio test for equality of CVs, *D*= 6.66, *n*_A0_=11,367, *n*_A-_=10,169). Further evidence comes from the shape of the distribution of mScarlet fluorescence. Populations with the A^−^ circuit exhibited mScarlet fluorescent distributions that are distinct and significantly narrower than populations with the A^0^ circuit (*p* = 2.2 × 10^−12^, Kolmogorov-Smirnov test: *D*=0.83; Figure 1D). Taken together, these results demonstrate that our A^−^ circuit successfully implements negative autoregulation.

### Adaptive evolution experiment of TetR toward novel inducer binding

To study how autoregulation affects the adaptive evolution of TetR towards responsiveness to a new environment we performed laboratory evolution experiments. We subjected TetR populations to random mutations and to selection for the ability to recognize a new inducer. Wild-type TetR recognizes the inducer tetracycline (Tc) or its non-toxic analog anhydrotetracycline (aTc) with high affinity^15,26^. As a new inducer (I_new_), we used the tetracycline analog 4-de(dimethylamino)-6-deoxy-6-demethyl-tetracycline (incyclinide), which is not a natural inducer of TetR. We chose this inducer, because previous studies have shown that directed evolution can increase TetR’s specificity to this inducer relative to tetracycline by more than 20,000-fold^26^. We performed all evolution experiments in four replicate populations for both A^−^ and A^0^ circuits.

For each replicate population and in every round (generation) of directed evolution, we introduced random mutations into the TetR gene by error-prone PCR, at a rate of approximately 1.5 nucleotide substitutions per TetR-coding gene. This rate corresponds to roughly 0.8 amino acid changes per TetR protein and generation (Materials and methods). Our population sizes ranged between approximately 10^5^ and 10^6^ individuals, such that the influence of genetic drift is negligible on the time scale of our experiment.

In each generation, we employed fluorescent activated cell sorting (FACS) to select cells for survival in a two-step process, which we refer to as negative and positive selection. During negative selection, we did not expose cells to the inducer I_new_ and allowed only cells to survive whose green fluorescence ranked within the lowest 20% of the uninduced population. This step is important, because it ensures the survival of cells in which TetR effectively represses GFP expression in the absence of an inducer. It prevents mutations that completely abolish TetR’s repressor activity from accumulating in a population, because such TetR mutants would show high green fluorescence. Following this negative selection, we proceeded with positive selection, which selects for TetR variants that can respond to the inducer I_new_. Specifically, we exposed cells to I_new_ and allowed only cells in the population’s top 5% of GFP fluorescence intensity to survive (Materials and methods). Together, this two-step scheme directly selects for the responsiveness of a circuit —requiring both repression in the OFF state and derepression in the ON state—while simultaneously maintaining the regulatory architecture itself. For each replicate population and for both A^−^ and A^0^ circuits, we repeated this sequence of mutagenesis and negative/positive selection for six consecutive generations.

### TetR responsiveness to the novel inducer increases during evolution

At the end of the experiment, we analyzed how TetR’s responsiveness to the inducer I_new_ had changed. GFP fluorescence serves as our quantitative readout of environmental responsiveness to the inducer. The fluorescence intensity of A^−^populations in the presence of I_new_ remained significantly higher throughout the evolution experiment (*p* < 10^−5^, generalized linear model; *n* = 4 independently evolving populations per circuit, Figure S25). A likely reason is the lower expression level of TetR in A⁻ populations, as reflected in their lower mScarlet fluorescence (Figure 1B). To allow us to compare changes in fluorescence intensity between A^−^ and A^0^ populations, it was thus necessary to normalize green fluorescence intensity. Specifically, for each circuit and in every generation, we subtracted the GFP fluorescence intensity of the uninduced TetR population from the fluorescence intensity of the population induced by I_new_ at that generation. We then divided the resulting quantity by the difference between the maximum fluorescence intensity of the population after induction with the inducer I_new_ (measured in generation six) and the fluorescence intensity of the uninduced state. This normalization scales fluorescence between 0 (uninduced state) and 1 (maximum fluorescence intensity observed at generation six). We refer to this quantity as the strength of induction by I_new_, or (for brevity) simply as induction. This quantity enables direct comparison of TetR responsiveness between A⁻ and A□ populations, despite their differing TetR expression levels.

The strength of induction by I_new_ increased over time during experimental evolution (Figure 2A), showing that TetR becomes increasingly responsive to the new inducer. Importantly, during the first three generations induction by I_new_ increased significantly more rapidly in the A^−^ population (generalized linear model, *p* = 0.0016, 0.0015, and 0.032, for generations 1-3, Figure 2A). After the third generation, it increased equally rapidly in both populations (*p*=0.050, 0.06, and 0.61, for generations 4-6, Figure 2A). Thus, not only did TetR become more responsive to I_new_, autoregulation facilitated its adaptive evolution early during the experiment. A closer examination of the repression of the GFP target gene with and without the inducer reveals why. It also demonstrates a key difference between A^0^ and A^−^ populations. In the absence of the inducer, A-populations evolve increased repression of the target gene compared to A^0^ population, which evolve decreased repression of the target gene (Figure 2B). Both populations, however, evolve higher derepression of the target gene in the presence of the inducer (I_new_) (Figure 2C). The differences between A^−^ and A^0^ populations hints that alleles rising to high frequency in A⁻ populations may possess different properties from those favored in A□ populations. To understand why the circuits evolve in different ways, we next studied the two main mechanistic aspects of switch behavior: TetR’s interaction with the inducer, which governs activation of the ON state, and TetR’s interaction with DNA, which maintains repression in the OFF state.

**Figure 2.**
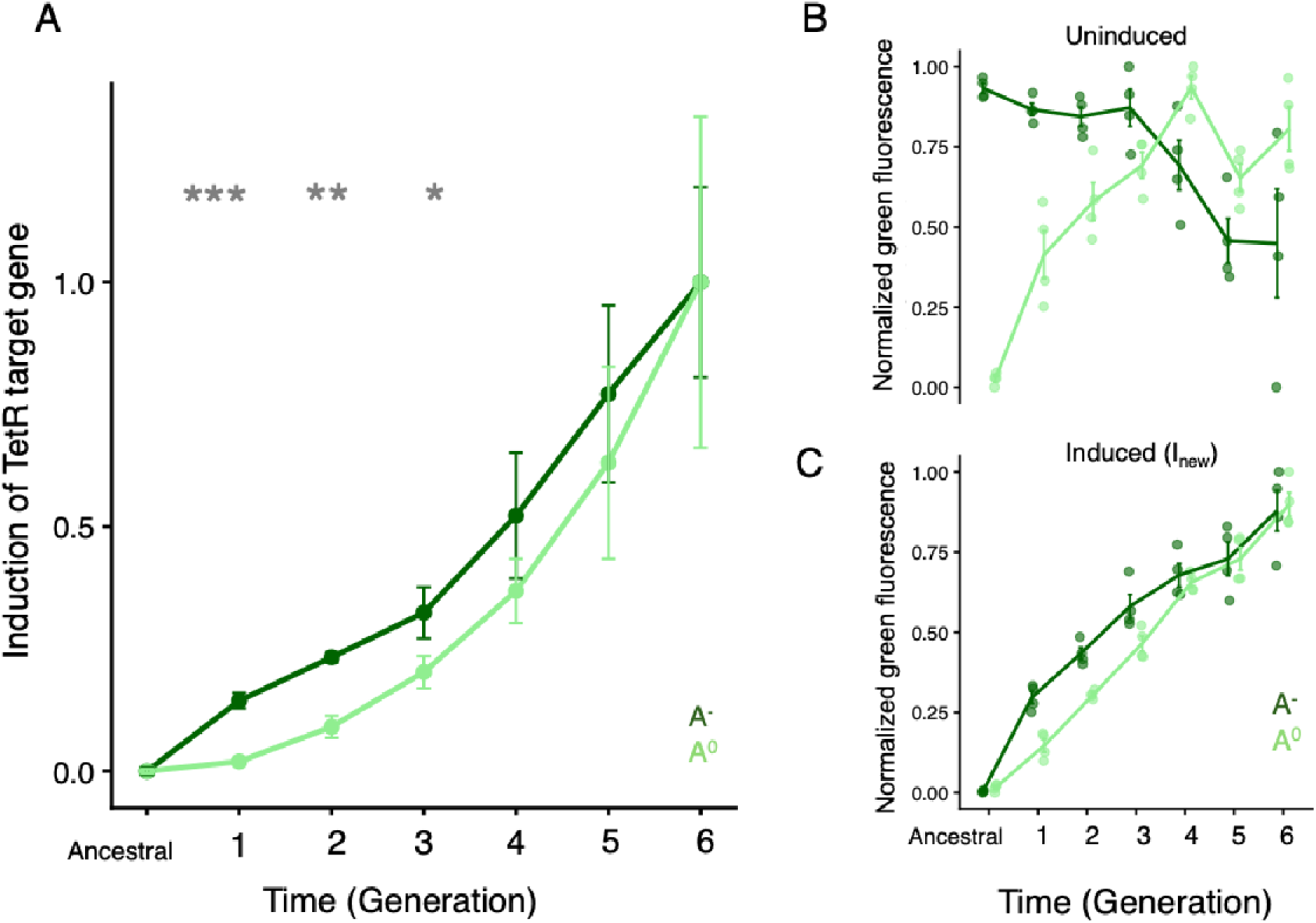
Evolution of TetR induction in circuits with **(A⁻)** and without (AT) negative autoregulation. A) Mean induction ratio of the TetR-regulated target gene by I_new_ across evolutionary rounds. The induction ratio is defined as the target gene expression (green fluorescence) in the presence of I_new_ divided by expression in its absence. Values were min–max normalized such that the ancestral (WT) induction ratio is set to 0 and the maximum observed induction (generation 6) is set to 1. This normalization enables direct comparison across populations that evolve in parallel.To assess differences in TetR induction across generations and populations (A⁻ vs. A□), we fitted a generalized linear model (GLM) assuming a Gaussian distribution. The model included time (generation of directed evolution) and replicate population as factors, along with their interaction. Statistical significance was determined based on GLM-derived p-values. For each round, the effect size (□, difference in normalized induction), its standard error, the test statistic (*t*), and two-sided *p*-value were obtained from the GLM (e.g., round I: □ = 0.13 ± 0.02, *t* = 5.40, *p* = 0.0017; round VI: □ 0, *t* = 0.53, *p* = 0.61). Full statistical results for all rounds are provided in Supplementary Table 9. Significance levels are indicated as follows: ∗ (*p* < 0.05), ∗∗ (*p* < 0.01), and ∗∗∗ (*p* < 0.001). B, C) Normalized green fluorescence intensity across evolutionary generations for both circuits under uninduced (panel B) and induced (by I_new_) conditions (panel C). In the uninduced state (panel B), TetR binds tightly to the GFP operator and strongly repress expression, which is indeed reflected by low absolute GFP levels in the both circuits (see Fig. 1C). The elevated values observed for the A⁻ population at generation 0 do not indicate high GFP expression but arise from the normalization applied across generations. Thus, the dark-green curve in panel B should be interpreted as the relative change in fluorescence across generations rather than the absolute GFP level in the absence of the inducer. In all panels, A⁻ and A□ populations are shown in dark and light green, respectively. Error bars represent the standard error of the mean (SEM) across replicate populations.

### Negative autoregulation favors TetR variants that interact more strongly with inducer

To study the first aspect of switch behavior—strong inducer responsiveness—we linked phenotypic evolution of this response to genotypic evolution in TetR. Specifically, we examined the evolutionary dynamics of genetic variants in both A⁻ and A□ populations by sequencing the TetR coding region of each biological replicate at every generation using single-molecule real-time (SMRT) sequencing (Materials and Methods; Supplementary Methods 4 and 5). On average, we obtained 36,689 high-quality TetR nucleotide sequences per replicate population and generation (Phred score per position > 30; Table S7). We then identified the ten alleles in A⁻ and A□ populations with the largest average frequency changes at the end of the experiment across replicates (Supplementary Figure S26). These alleles most likely harbor the mutations driving adaptive evolution. Together, they introduced changes at 14 TetR amino acid positions, six within the inducer-binding domain and eight within the DNA-binding domain.

Among these candidates, only a few alleles exhibited consistent frequency increases across all replicates and reached moderate frequencies (>0.1) by the final round of evolution (Figure 3A, Supplementary Figure S26). In the A⁻ populations, R105W was the most prominent such allele, rising to the highest frequency in every replicate, and reaching an average final frequency of 0.6 ± 0.34 across replicates. In the A□ populations, P106L was the only allele whose frequency consistently increased across all replicates (Table S8). It reached an average final frequency of 0.95 ± 0.05 across the replicates (Figure 3A). This pattern suggests that different regulatory architectures (negative autoregulation in A⁻ and its absence in A□ populations) might favor the selection of distinct genotypes in our experiments. We focused our subsequent functional characterization on R105W (from A⁻ populations) and on P106L (from A□ populations), because they are the strongest candidates for adaptive changes in TetR in the two circuit types.

**Figure 3.**
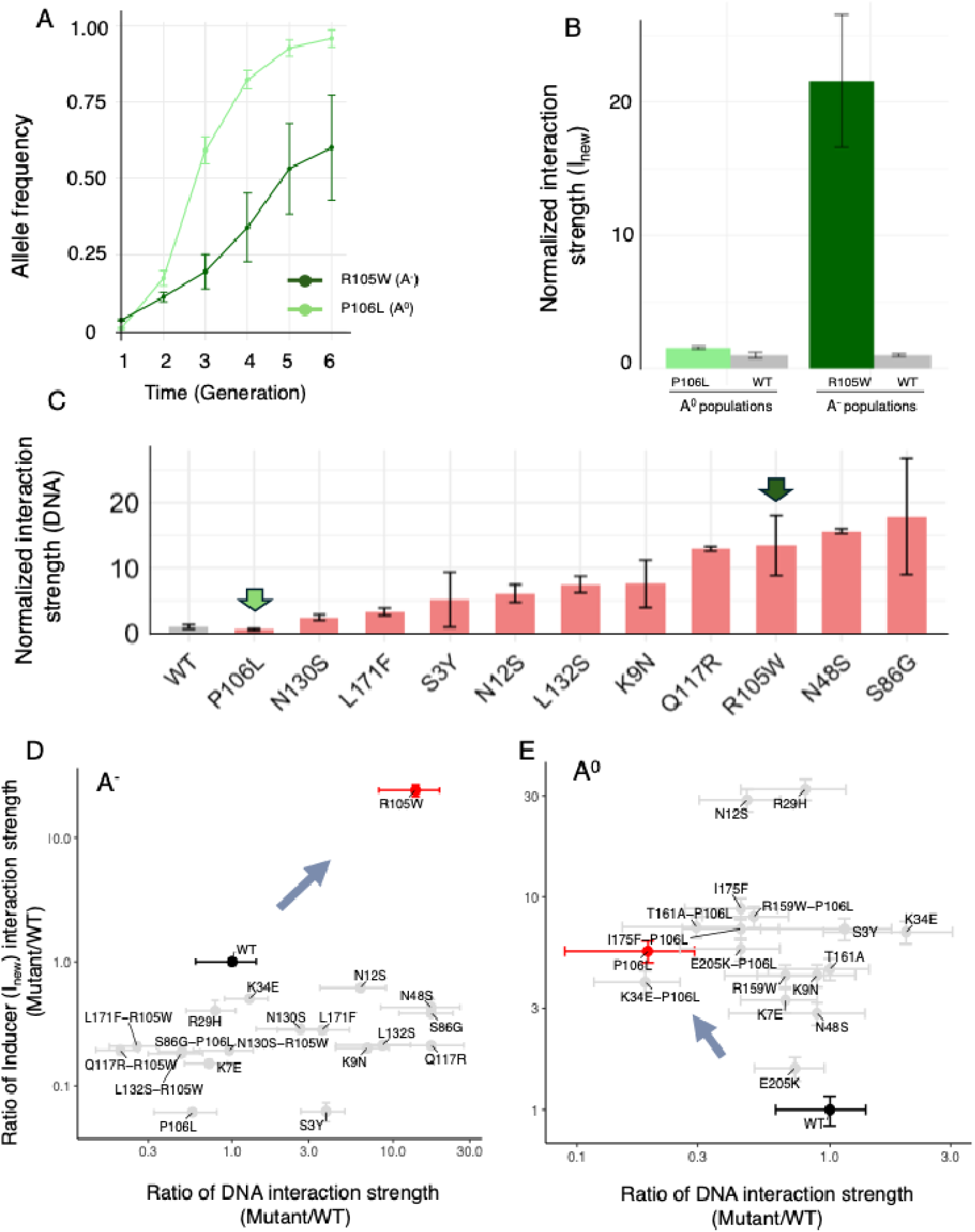
TetR variants with distinct properties are selected in NAR and NoNAR populations. A) The panel shows the frequency of alleles R105W and P106L (vertical axis) plotted against time (generations of evolution). These alleles reach the highest frequencies among all alleles in the A⁻ and A□ populations, respectively. B) Normalized interaction strength of TetR variants with the inducer I_new_ for R105W (dark green) and P106L (light green). The interaction strength is normalized by the values of WT TetR in the corresponding circuit, so WT values in this panel are equal to 1 and are shown in gray. We inferred the inducer interaction strength by fitting a one-site binding equation to experimental data on reporter green fluorescence, and by estimating the interaction strength as the reciprocal of the dissociation constant (see Supplementary Methods 14). C) Normalized interaction strength (vertical axis) of TetR with DNA for TetR alleles specific to the A⁻ population (horizontal axis). We estimated interaction strength with DNA with the same procedure as in (B), but through reporter mScarlet fluorescence, which reflects how tightly TetR binds to the mScarlet promoter in the A⁻ circuits, and how this binding depends on I_new_ concentration. (Supplementary methods, section 14). The two variants that reached the highest frequencies in A⁻ (R105W) and A□ (P106L) populations are highlighted with arrows. D, E) Normalized inducer interaction strength (vertical axes) is plotted against normalized DNA interaction strength (horizontal axes) for TetR alleles that rose to high frequencies in A⁻ populations (panel D) and A^0^ populations (panel E). Each grey circle represents data from a TetR single mutant or a double mutant, with the ratio of interaction strength calculated relative to the wild-type (WT). Error bars denote standard deviations derived from measurements of inducer and DNA interaction strength in n=4 biological replicate populations. WT values are shown as black circles with corresponding error bars. Alleles R105A and P106L are highlighted in red in both panels, because they are prevalent alleles in A⁻ and A^0^ populations, respectively. Allele R105W exhibits both strong DNA interaction and strong inducer interaction relative to the WT (direction of arrow), distinguishing it from other alleles. Allele P106L and its double mutant combinations show strong inducer interaction but weak DNA interaction compared to the WT.

Both amino acids R105 and P106 are located within the inducer-binding pocket of TetR, and help to stabilize inducer binding through hydrophobic interactions^26^. In addition, the amino acid changes that occur at these positions are changes towards more hydrophic amino acids (Arginine (R) to Tryptophan (W) in R105W, and Proline (P) to Leucine (L) in P106L) (Figure S26). This suggests that increased inducer affinity helps to drive the frequency increase of the mutant alleles R105W and P106L.

To test this hypothesis and quantify the strength of interaction between TetR and inducer, we engineered variants R105W and P106L into ancestral TetR, and measured green fluorescence intensity following induction with I_new_. We conducted a titration assay by varying the concentration of I_new_ over a range from 10^−3^ to 10^2^ µM and recorded the resulting changes in green fluorescence (see Supplementary Methods 14). While this assay does not directly measure inducer binding affinity, it provides a functional readout of TetR–inducer interaction, which we refer to as the *inducer interaction strength*. To control for circuit-specific effects, we compared the interaction strength of each mutant to that of the corresponding WT TetR allele within the same circuit background.

Both alleles exhibited increased inducer interaction strength compared to the wild-type (WT) TetR, but with different magnitudes (Figure 3B). R105W, the allele that consistently rose to the highest frequency in A⁻ populations, showed a ∼25-fold increase in interaction with inducer relative to WT in the A⁻ circuit. In contrast, P106L, the leading allele in A□ populations, showed only a 1.5-fold increase relative to the WT in the A□ circuit. This stark difference suggests that the presence of negative autoregulation favors the selection of alleles with substantially stronger inducer interactions, such as R105W.

### Negative Autoregulation Promotes the Selection of TetR Variants with Enhanced DNA Interaction Strength

We next studied the second aspect of switch behavior, strong interaction of TetR with DNA. For an environmentally responsive molecular switch, strong DNA binding is as important as strong inducer binding, because it dictates the strength of repression in the absence of inducer. DNA binding and inducer binding are allosterically coupled in TetR. When TetR binds its inducer, a conformational change reduces its affinity for DNA, thereby relieving repression^15,16^. This coupling implies that TetR alleles with altered inducer-binding strength can also show altered DNA binding.

In our circuits, the gene encoding mScarlet is immediately downstream of the TetR gene, and its expression is modulated by TetR’s autoregulatory binding to the *tetO* operator. Thus, changes in mScarlet fluorescence reflect changes in TetR’s interactions with DNA. We refer to this responsiveness as DNA interaction strength, a proxy for DNA-binding affinity. To test whether R105W or P106L alter TetR–DNA interactions, we performed titration assays across a wide range of inducer concentrations (10^−3^ to 10^2^ µM) and quantified the sensitivity of mScarlet fluorescence (see Supplementary Methods 14).

We observed dramatic differences in the DNA interaction strengths of R105W (the prevalent allele in the A⁻ population), and P106L (the prevalent allele in A□ populations). While R105W was among the top three alleles with the highest DNA interaction strength (Figure 3C), P106L interacted most weakly with DNA among all alleles we tested. We also found that DNA interaction strength and inducer interaction strengths are largely independent of each other – they are not significantly correlated among the TetR alleles we examined (*R*=0.15, *p*=0.51, Spearman’s rank correlation, *n*=20 alleles). Therefore, the higher DNA interaction strength of R105W compared to P106L is not a by-product of increased inducer interaction strength.

We next compared inducer and DNA interaction strengths of the top 10 high frequency variants in both A□ and A⁻ populations. To permit comparison of these quantities among circuit types, we normalized them to that of WT TetR. Figure 3 highlights these quantities for the alleles R105W (Figure 3D, A^−^ populations) and P106L (Figure 3E, A^0^ population) in red, along with their respective WT controls (black). These data reveal that selection favored different kinds of interactions in circuits with autoregulation. Specifically, the prevalent variant in A^−^ populations, R105W, stands out because it interacts strongly with the inducer I_new_ and with DNA. In contrast, the prevalent variant in AO populations, P106L, interacts strongly with the inducer I_new_, but not with DNA.

Biologically, a TetR variant that interacts strongly with DNA according to our assay can effectively repress its target promoter. When this variant also binds inducer strongly, the circuit responds sharply to environmental changes. Such a repression enables strong repression when inducer is absent, and strong derepression when inducer is present. The combination of the two entails strong induction by I_new_, as we observed in our experiments (Figure 3A). R105W exemplifies this strong responsiveness to the environment. These favorable properties explain the high frequency of this variant in A^−^populations (∼0.6 ± 0.34).

Despite these properties, the frequency of this variant remained extremely low in A□ populations (∼0.0020 ± 0.0018; Table S8). Conversely, in A□ populations the variant P106L (which only achieved a modest frequency of ∼0.16 ± 0.20 in A⁻ populations) rose to the highest frequency (∼0.95 ± 0.05). These differences underscore that mutations that increase DNA interaction strength like R105W are only favored under negative autoregulation. To understand why, we developed a simple biophysical model of TetR repression to study the evolutionary dynamics of TetR.

### A mechanistic model based on the biophysics of inducer binding and negative autoregulation

The model tracks the evolution of TetR’s key biophysical properties, inducer-binding affinity and DNA-binding affinity, under selection for high inducible expression of a target gene. The model represents populations of individual cells (Supplementary Results 4). In each cell, TetR is characterized by two parameters: DNA-binding affinity (*A_d_*), defined as the ratio of TetR bound to the promoter relative to free TetR, and inducer-binding affinity (*A_ind_*), the ratio of inducer-bound TetR to free TetR (Figure 4A). TetR expression is either constant (in the A□ circuit) or repressed by negative autoregulation based on equilibrium thermodynamics (in A^−^ circuits, Supplementary Results 4). We modeled mutational change during the evolution of *A_d_* and *A_ind_* by independently adding values to these parameters that are drawn from normal distribution centered at zero, with a standard deviation of 10% of the current parameter value. After each round of mutation, we allowed the top 5% of cells with the highest induced GFP fluorescence to survive to seed the next generation.

**Figure 4.**
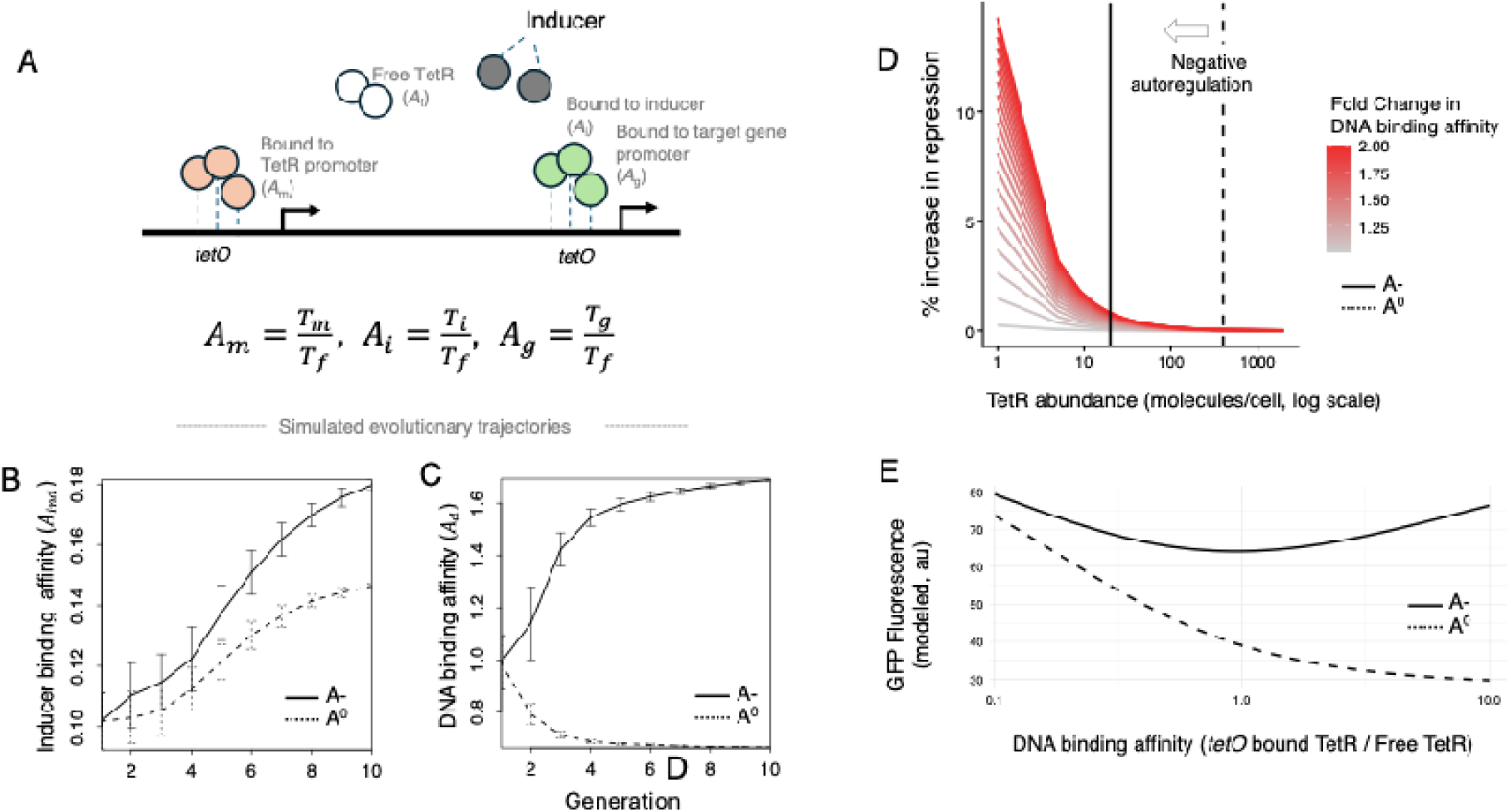
A mechanistic model of how TetR interacts with inducer and DNA helps to explain the evolutionary dynamics of TetR variants in **A^0^ and A^−^** populations. **A)** Schematic representation of the TetR regulatory system and its binding states. TetR can exist in one of several forms: bound to the TetR promoter (T_m_), bound to the target gene promoter (T_g_), bound to the inducer (T_i_), or free (T_f_). The ratios *A*_m_=T_m_/T_f_, *A*_i_=T_i_/T_f_, and *A*_g_=T_g_/T_f_ quantify effective interaction strengths of TetR with its respective partners, and form the basis of the mechanistic model. This model allows quantification of DNA and inducer binding properties in circuit-independent units. B, C) Simulated evolutionary trajectories for inducer binding affinity (*A_ind_*, panel B), and DNA binding affinity (*A_d_*, panel C) and during ten rounds (generations) of mutation and selection, starting from the initial conditions *A_d_*=1, and *A_ind_*=0.1 (see Methods for details of evolutionary simulations). Each line represents the evolution of mean within-population TetR affinity over time (generations, horizontal axis; error bars: 1 standard deviation) from a simulated A⁻ (solid line) and A□ (dashed line) populations. The panel shows that A⁻ populations evolve higher DNA binding and stronger inducer binding than A□ populations. D) Percentage increase in target gene repression (in the absence of inducer) relative to the WT TetR, as a function of TetR abundance (horizontal axis), and for different modeled alleles with different DNA binding affinity relative to the WT (color legend, one curve per allele). Each curve corresponds to a modeled TetR allele whose DNA binding affinity (A_d_) is higher than that of the TetR WT (*A*_d_=1) by the fold increase shown in the color legend. We quantify the increase in target gene repression as the percentage decrease in modeled GFP fluorescence relative to the WT. We note that in the absence of inducer DNA binding affinity (*A*_d_) is the sole parameter that affects repression. The panel shows that as the abundance of TetR increases, the percentage of repression caused by an allele with increased DNA binding affinity relative to the WT decreases. The reason is that for alleles with increasing binding affinity the TetR target gene promoter is more likely to be saturated at a given TetR concentration. What negative autoregulation achieves is depicted by the arrow and the shifted position of the solid line relative to the dashed line: It reduces TetR abundance and thus saturation of the target gene promoter, rendering repression more sensitive to increased DNA binding. E) Predicted relationship between DNA binding affinity (x-axis; measured as the ratio of TetR bound to DNA versus free TetR) and induced target gene expression (GFP fluorescence, y-axis) in the modeled A⁻ (solid line) and A□ (dashed line) populations. The model predicts that without negative autoregulation (in A□ populations), target gene expression declines as TetR DNA binding affinity increases. In contrast, with negative autoregulation (in A⁻ populations), target gene expression increases again at high DNA binding affinity due to autoregulation-driven reduction in TetR abundance.

In studying our model’s predictions, we focused mostly on the biologically realistic regime where DNA-binding affinity (*A*_d_) is substantially greater than inducer-binding affinity (*A*_ind_), i.e., *A*_d_ ≫ *A*_ind_. This assumption is supported by empirical measurements of the tetracycline-TetR system. In the absence of tetracycline or its derivatives, TetR binds its cognate operator, *tetO*, with extremely high affinity, with dissociation constants in the subnanomolar range (∼10^−10^ M)^24^. Strongly binding inducers such as anhydrotetracycline (aTc) also bind TetR with high affinity (∼10^−12^ M)^24,27^, and this binding induces a conformational change that disrupts TetR’s DNA-binding interface, reducing its DNA-binding affinity to approximately 10^−6^ M for the aTc-bound form^24^. This implies that strong inducer binding shifts TetR into a state where its ability to bind DNA is markedly weakened. However, at the beginning of evolution towards interaction with a new inducer I_new_, TetR alleles likely still bind poorly to the new inducer, such that DNA binding is prevalent in its influence on gene expression. In addition to simulations within the most realistic regime of strong DNA binding, we also performed simulations across a broad range of initial values of *A_d_* and *A_ind_* to assess how sensitive our results are to the choice of starting parameters (Figure S29).

The model predicts that induced target gene expression increases over time in both A⁻ and A□ populations, mirroring the experimental rise in GFP derepression (Supplementary Results 4; Supplementary Table S6; Figure S28). Importantly, when we initiated simulations with a TetR variant that binds DNA much more strongly than inducer, as is the case for the initial TetR wild-type in our experiments, the model’s evolutionary dynamics closely recapitulated the experiments’: Target gene induction increases in both A^−^ and A□ populations, but does so initially faster in A^−^ populations, before the two populations converge in the strength of induction (Figure 2C, Figure S28).

The model also recapitulates the key experimental differences between A□ and A⁻ populations in the evolution of *A*_d_ and *A*_ind_. In both kinds of populations, the model predicts that TetR molecules consistently evolve high inducer-binding affinity *A*_ind_ (Figure 4B), as our experiment do (Figure 3B). However, the model also predicts that only A^−^ populations favor TetR molecules with high inducer-binding affinity *A*_ind_ and high DNA-binding affinity *A*_d_ (Figure 4C). This difference validates our experimental observations, where the prevalent allele in the A⁻ population, R105W, exhibits stronger DNA interaction (Figure 3D), whereas the prevalent allele in the A□ population, P106L, displays weaker DNA interactions (Figure 3E).

This difference between A□ and A⁻ populations arises because negative autoregulation fundamentally reshapes how DNA binding strength affects the induction of TetR’s target gene. In the absence of inducer, autoregulation lowers the steady-state abundance of TetR in A⁻ circuits relative to A□ circuits, which renders the repression of the target gene more sensitive to increases in TetR’s DNA binding affinity *A*_d_. Specifically, at lower TetR abundance, even modest increases in TetR’s DNA-binding affinity can produce large increases in OFF-state repression (Figure 4D). In contrast, A□ circuits operate at much higher TetR abundance, where the same increase in *A*_d_ yields a lower repression gain. Upon addition of inducer, A⁻ circuits then gain a second advantage: because autoregulation keeps TetR levels low, TetR variants with high inducer affinity can sequester a larger fraction of the TetR pool in the inducer-bound state, producing stronger derepression and a greater strength of induction (Figure 4E). Thus, variants in which both *A*_d_ and *A*_ind_ are increased simultaneously tighten OFF-state repression and amplify ON-state induction—but only in the presence of negative autoregulation.This explains why strongly DNA-binding mutations are favored in A⁻ but not in A□ populations.

## Discussion

We find that negative autoregulation alters the fitness landscape of TetR, such that TetR alleles that would be deleterious in the absence of negative autoregulation, can become highly adaptive in its presence. This is best illustrated by allele R105W. This allele was not beneficial in A□ populations (without negative autoregulation), but it was highly beneficial in A⁻ populations (with negative autoregulation), where it rose to highest frequency in all replicate populations. This allele exhibits the strongest interaction with both inducer and DNA among all TetR variants we tested, and allowed A□ circuits to respond most strongly to the environment: Specifically, it allowed strong repression in the absence of inducer, and strong derepression in the presence of inducer—precisely the properties sought in an environmentally responsive molecular switch.

The selection of this allele is a direct consequence of the regulatory architecture we study. In A⁻ circuits, strong DNA binding has consequences different from A□ circuits. Specifically, stronger DNA binding lowers TetR abundance through negative autoregulation, which has two effects. First, in the absence of inducer, the promoter is no longer saturated by TetR, so an increase in DNA-binding affinity by mutation of TetR produces a disproportionately large increase in repression (Figure 4D). This renders such mutations more beneficial in the OFF state. Second, in the presence of inducer, a smaller TetR pool due to autoregulation means that fewer TetR molecules remain bound to the promoter, making derepression easier to achieve. In this way, negative autoregulation enables the selection of alleles like R105W, which combines strong DNA and inducer interactions to maximize environmental responsiveness, but which would otherwise remain unfavorable.

The pattern we observed for R105W is an example of mutual exclusivity, where the same allele is adaptive under one regulatory architecture but maladaptive under another. Other alleles show a similar pattern. Specifically, we examined all 2314 alleles detected in both A⁻ and A□ populations by the sixth round of evolution. For each allele, we quantified its mean frequency in both A^−^ and A^0^ populations (Figure S30). Alleles that rose to high frequency in one population tend to remain at low frequency in the other, resulting in a highly significant orthogonal pattern of allele frequencies (*p*-values < 10^−16^, Fisher’s exact test; Figure S31). This observation suggests that regulatory architecture can make multiple mutations adaptive in one context but deleterious in another.

The most obvious limitation of our work is that it relies on a synthetic system, in which TetR is embedded into a minimal circuit that does not occur in nature. A synthetic system eliminates many of the complexities that govern transcriptional regulation in living cells, including crosstalk between regulators and multi-layered signaling cascades^5,28,29^. At the same time, it also allows us to isolate one central feature—negative autoregulation—and evaluate its evolutionary consequences without confounding effects.

A second limitation is that our measurements of interaction strength are indirect, inferred from reporter fluorescence rather than obtained through direct biophysical assays. However, because repression and derepression occur at the transcriptional level and are directly reflected in the expression levels of our target genes, fluorescence provides a meaningful proxy for the switching behavior we study. This approach allowed us to compare evolutionary trajectories across many variants in a high-throughput manner. Nevertheless, future work that measures TetR–DNA and TetR–inducer interactions directly, for example using electrophoretic mobility shift assays^30^, surface plasmon resonance^31,32^, microscale thermophoresis^33^, isothermal titration calorimetry^34,35^, yeast surface display, or phage display^36^, may help to disentangle direct binding from indirect effects, for example of protein maturation and stability.

Lastly, the form of selection we applied in our experiments is threshold-like. It favors TetR variants that exceed a fixed GFP fluorescence cutoff. Although this selection is artificial, it has biological parallels. Thresholds are important in many natural phenomena such as antibiotic resistance^37^ and quorum sensing^38^. For example, in competence development of *Bacillus subtilis*, the master regulator ComK must accumulate beyond a threshold level to enable cells to take up DNA^39^. And in morphogen-driven developmental decisions, gradients of signaling molecules, such as Bicoid in *Drosophila* or Sonic Hedgehog in the vertebrate neural tube, are transformed into discrete cell fates based on threshold amounts of these molecules^40,41^. Thus, although our experimental selection scheme is stylized, it captures an important mode of natural selection that is directly relevant to the evolution of environmentally responsive molecular switches.

Negative autoregulation has multiple well-established physiological effects. It can accelerate a circuit’s response time, reduce noise, and stabilize gene expression^22,23,42^. Our work shows that it can also affect adaptive evolution. It changes which mutations in a regulator are adaptive. It alters how DNA and inducer interactions translate into the repression and derepression of a circuit’s target gene. And it helps to evolve circuits that respond strongly to environmental change.

## Acknowledgments

We would like to acknowledge financial support by Swiss National Science Foundation grants 31003A_172887 and 310030_208174. We thank the Functional Genomics Center as well as the Cytometry Facility at the University of Zurich for technical support. PD acknowledges Federal Ministry of Education in Germany (BMBF), and research grant under the funding code 01EK2203A.

## Materials and Methods

### Strains and Plasmids

We used *E. coli* strain DH5□ for all cloning steps and experiments. We used SIG10 HIGH Electrocompetent cells (Sigma-Aldrich CMC0003) for library cloning and directed evolution experiments. All strains and vectors are listed in Supplementary Tables S1 and S2.

### Directed evolution using fluorescence-activated cell sorting (FACS)

We constructed plasmids with or without negative autoregulation starting from a plasmid named pDC-NAR-MCS, which we had synthesized by Twist Bioscience (San Francisco, CA). The plasmid encodes both mScarlet and sfGFP fluorescent proteins, driven by a modified BBa_J23100 promoter. The promoter is repressed by TetR via a tetO2 binding site placed downstream of the −10 region. We generated pDC-NoNAR-MCS by replacing the tetO2 sequence with a scrambled variant (5’-AGAACTAGGTATCCGCTAT-3’) upstream of mScarlet. We then cloned the TetR gene upstream of mScarlet in pDC-TetR-NAR and pDC-TetR-NoNAR. Below, we use the acronym ‘A^−^’ to refer to the plasmid construct with negative autoregulation (pDC-TetR-NAR) and ‘A^0^’ to refer to the plasmid construct without negative autoregulation (pDC-TetR-NoNAR).

We confirmed the desired outcome of cloning procedures by restriction digestion and sequencing (Supplementary Methods 4, and Supplementary Figure S1). We introduced random mutations into TetR using a mutagenic polymerase chain reaction (PCR), with a reaction mixture containing template plasmid, dNTPs, Taq polymerase, and MnCl2. After PCR amplification, we ligated the linearized TetR inserts into the vector backbone and electroporated the ligation products. Mutagenesis resulted in approximately 1.5 nucleotide mutations per TetR molecule, as determined by SMRT sequencing (Supplementary Methods 5, Supplementary Table S4). We performed FACS on a BD FACSAriaIII cell sorter, setting PMT voltages to 478V (FSC), 282V (SSC), 480V (FITC), and 490V (PE-Texas Red). We excluded debris using a threshold of 1000 on FSC-H and SSC-H.

Directed evolution was performed using fluorescence-activated cell sorting (FACS) in two sequential selection steps per round. First, negative selection was applied to eliminate clones that showed high GFP expression in the absence of inducer—i.e., those with constitutive derepression. For this, transformants were grown overnight in M9 medium *without* inducer, and the lowest 20% of the population based on green fluorescence (FITC channel) was collected and carried forward. This step ensures that only functional repressors (i.e., TetR variants capable of DNA binding) were retained. Second, positive selection was applied to enrich for mutants responsive to the inducer. Cells from the negative selection step were grown overnight in M9 medium supplemented with 42 ng/ml of the novel inducer incyclinide. We then sorted the top 5% of green fluorescent cells, corresponding to high GFP expression upon derepression, using a BD FACSAriaIII (PMT voltages: 478V FSC, 282V SSC, 480V FITC, 490V PE-Texas Red). Debris was excluded using a 1000 threshold on FSC-H and SSC-H. Each replicate population underwent this two-step selection process across six evolutionary rounds. Full FACS protocols and gating strategies are detailed in Supplementary Methods 6–7.

After each generation of evolution, we analyzed the phenotype of evolving populations using a BD Biosciences LSR Fortessa flow cytometer, recording 10,000 cells per sample. We measured GFP fluorescence in the FITC channel and mScarlet fluorescence in the red channel. We analyzed the resulting data using the R package flowCore (Supplementary Methods 8). We sequenced the TetR coding sequence from plasmid libraries isolated after each generation of evolution using single-molecule real-time (SMRT) sequencing (Pacific Biosciences, PacBio). For multiplexed sequencing, we generated barcoded libraries according to PacBio’s instructions. To this end, we performed two generations of PCR. In the first generation, we amplified the TetR coding region using primers with a “universal” sequence. In the second generation, we used barcoded primers to differentiate the samples. After assessing PCR product purity, we pooled and purified the products before sequencing them on a PacBio SequelII SMRT cell (Supplementary Methods 9). Additional details on library construction and sort-seq procedures are provided in the Supplementary material.

### Validation of TetR mutants using binding assays

To generate single and double mutants in A^−^ and A^0^ constructs, we amplified the tetR gene from A^−^ (pDC-TetR-NAR) and A^0^ (pDC-TetR-NoNAR) plasmids, introducing the desired mutations via PCR. We then purified the PCR products and performed Gibson assembly to recombine the mutant fragments with the digested vector backbones. After transformation of the assembled vector into electrocompetent SIG10 cells we confirmed correct construction by DNA sequencing (Supplementary Methods 11).

We cultured individual TetR mutants overnight at 37°C in M9 medium supplemented with 0.4% glucose, 0.2% casamino acid, 2 mM magnesium sulfate, 0.1 mM calcium chloride, 50 µg/ml kanamycin, and either 1 µl/ml anhydrotetracycline (aTc) or 2 µl/ml incyclinide. We tested each of these condition in triplicate with three biological replicates of each mutant. Following growth, we washed the cells, resuspended them in PBS, diluted this resuspension and transferred the final dilution to 96-well plates for flow cytometry analysis. We measured GFP and mScarlet fluorescence using a BD Biosciences FACSymphony cell analyzer, analyzing 10,000 cells per replicate (Supplementary Methods 13, Supplementary Figures S8-S15).

We measured the interaction strength between TetR mutants and inducers anhydrotetracycline (aTc) and incyclinide and between TetR mutants and DNA in response to incyclinide using fluorescence-based assays in 96-well plates with a plate reader (Spark, Tecan). To this end, we grew mutants overnight at 37°C in M9 medium supplemented with 0.4% glucose, 0.2% casamino acid, 2 mM magnesium sulfate, 0.1 mM calcium chloride, and 50 µg/ml kanamycin. We assayed interaction strength to each inducer and DNA in triplicate with three biological replicates per mutant. For the inducer binding assay, we prepared two-fold serial dilutions of aTc and incyclinide across 16 concentrations in M9 medium supplemented as just described. For the DNA interaction assay, we prepared wells with 100 µM incyclinide and then performed a series of two-fold serial dilutions. We added 20 µl of the overnight culture to each well containing 180 µl of the respective diluted inducer solution in this M9 medium. We measured GFP fluorescence at different inducer concentrations to quantify interaction strength for each inducer. For estimating the interaction strength between TetR mutants and DNA, we measured mScarlet fluorescence in response to the new inducer incyclinide. We used mScarlet fluorescence to evaluate DNA interaction strength, because it serves as a proxy for TetR expression, which directly depends on the amount of free TetR available to bind DNA. We normalized both GFP and mScarlet fluorescence to optical density at 600nm (OD_600_) and performed min-max normalization for each replicate for all mutants to ensure comparability across samples. We plotted fluorescence data against inducer concentrations and calculated dissociation constants, K_D_, between each mutant and the inducer or between each mutant and DNA using a one-site specific binding equation (Supplementary Methods 14, Supplementary Figures S16-S24).

### Data analysis and simulations

Primary data analysis was performed by the Functional Genomics Center Zurich. This included generating consensus reads from sub-reads and demultiplexing the SMRT sequencing data using the SMRTAnalysis v2.3 package (12). We mapped reads from each resulting data file to the ancestral TetR sequence using custom R scripts with SAMtools (13). We used only mapped reads spanning the entire TetR coding region with an average Phred quality score above 20 for further analysis. After excluding sequences lacking a start or stop codon, we were left with at least 10^4^ sequences from each replicate population for further analysis. We wrote custom Python scripts (Python 2.7.12) to identify point mutations and convert them to amino acid sequence changes. We then used custom-made R (v4.2.1) scripts to calculate the frequencies of different amino acid variants in each replicate population and simulate adaptive evolution.

We simulated the evolution of TetR’s DNA-binding affinity (*A*_d_) and inducer-binding affinity (*A*_ind_) using individual-based simulations, in which each cell is represented by these two parameters. Each population consisted of 10³ cells carrying either an A□ or A⁻ circuit. In every generation, mutations were introduced by perturbing *A*_d_ and *A*_ind_ with random effects drawn from a normal distribution centered at zero with a standard deviation of 10% of the current parameter value. Cells were then evaluated based on their induced GFP fluorescence (i.e., fluorescence in the presence of inducer relative to the uninduced state). we derived equations that describe how induced GFP fluorescence depends on *A*_d_ and *A*_ind_ under the two circuit architectures: constant expression in A□ and negative autoregulation in A⁻ (Supplementary Result 4). The top 5% cells with the highest modeled induced GFP fluorescence were allowed to contribute to the next generation. From these survivors, the mean and standard deviation of *A*_d_ and *A*_ind_ defined the parameter distributions from which the next generation was drawn. Population size was held constant across generations. This process was repeated for 10 generations to track evolutionary changes in fluorescence and binding affinities (see Supplementary Results 4 for full model description and Table S6 for parameter values). Scripts for sequence analysis and evolutionary simulations are available at: https://github.com/dasmeh/NegAutoReg

